# Fluoroquinolone-Specific Resistance Trajectories in *E. coli* and their Dependence on the SOS-Response

**DOI:** 10.1101/2024.06.06.597756

**Authors:** Lisa Teichmann, Sam Luitwieler, Johan Bengtsson-Palme, Benno ter Kuile

**Affiliations:** University of Amsterdam, Swammerdam Institute of Life Sciences, Molecular Biology and Microbial Food Safety, Amsterdam, The Netherlands; Chalmers University of Technology, Department of Life Sciences, SciLifeLab, Division of Systems and Synthetic Biology, Gothenburg, Sweden; University of Gothenburg, Institute of Biomedicine, Department of Infectious Diseases, Gothenburg, Sweden; Centre for Antibiotic Resistance Research (CARe) in Gothenburg, Sweden

**Keywords:** Fluoroquinolone resistance, *de novo* resistance, SOS response, resistance mutations

## Abstract

Fluoroquinolones are essential for treating bacterial infections in both human and veterinary medicine. This study investigates the mechanisms behind acquired resistance to fluoroquinolones with a specific focus on the SOS response - a critical cellular pathway activated by DNA damage. Utilizing an experimental evolution approach, we exposed *Escherichia coli* to four fluoroquinolones and monitored the adaptation process. A *recA* knock-out mutant deficient in the SOS response was used as biological control. The emergence of resistance was accompanied by numerous DNA mutations, consisting of some observed often and others that infrequently appeared. Our findings indicate that the development of resistance depends in varying degrees on the SOS response among the tested fluoroquinolones, with notable dissimilarities in clinical resistance development. Resistance developed slowest to ciprofloxacin, then levofloxacin, followed by enrofloxacin, and fastest to moxifloxacin. Genomic analysis revealed distinct mutation profiles in cultures exposed to the tested antimicrobials, emphasizing the unique adaptation strategies of bacteria. This research underscores the importance of recognizing the differences among fluoroquinolones in scientific research and clinical practice.

## 1. Introduction

Fluoroquinolones are broad-spectrum antibiotics commonly used to treat infections in both human and veterinary medicine (1,2). This class of antibiotics is listed within the Watch group, a classification devised by the World Health Organization (WHO) under the AWaRe framework established in 2017 (3). Antibiotics classified in this group are considered to have a heightened potential for resistance and are categorized as the highest priority critically important antimicrobials for human medicine. Despite their importance and our extensive knowledge of their mechanisms of action and corresponding bacterial resistance mechanisms, our understanding of how bacteria evolve to become resistant or adapt to fluoroquinolones remains limited.

The mode of action of fluoroquinolones revolves around targeting essential bacterial enzymes pivotal for DNA replication and repair, i.e. DNA gyrase (GyrA and GyrB) and DNA topoisomerase IV (ParC and ParE) (4). The bacterial resistance mechanisms include target site alterations, efflux pumps, and changes in membrane permeability (5). Since fluoroquinolones cause DNA damage and consequently have been shown to activate the bacterial SOS response, inhibiting the SOS response has been proposed as a potential approach to combat resistance evolution (6,7).

At the core of the SOS response lays the activation of the RecA protein, induced by its interaction with single-stranded DNA (ssDNA) (8,9). Once activated, RecA coordinates the self-cleavage of the SOS transcriptional repressor LexA (10), thereby initiating the transcriptional cascade of the SOS response. Knocking out RecA does not entirely abolish antibiotic-induced mutations in *Escherichia coli* upon beta-lactam exposure, indicating the existence of a LexA/RecA-independent pathway capable of triggering the SOS response that has yet to be fully understood (11).

Utilizing an experimental approach aimed at inducing the evolution of resistance, we sought to unravel the dynamics governing the adaptation of *E. coli* to four fluoroquinolone antibiotics – ciprofloxacin, levofloxacin, moxifloxacin, and enrofloxacin – using a SOS-deficient mutant as biological control. The selection of these fluoroquinolones was guided by their diverse chemical structures and widespread use in both veterinary and human healthcare settings, allowing for a broad exploration of the adaptive landscape. Yet, how and if they differ in inducing cellular changes that lead to *de novo* resistance is unknown. Through this study, we aimed to elucidate the genetic and phenotypic changes occurring in both the wild type and SOS-knockout strain, thus highlighting the interplay between the induction of the SOS response and the evolution of resistance. Our research specifically focused on the role of the SOS response in adaptation to each fluoroquinolone and the distinct adaptation patterns associated with each antibiotic.

Our results suggest significant disparities in the adaptational response of *E. coli*, as well as its dependency on the SOS response, to various fluoroquinolones. Specifically, we observed that the knockout mutant and the wild-type strain exhibited varying rates of clinical resistance development, with the fastest adaptation occurring after exposure to moxifloxacin and the slowest to ciprofloxacin. Moreover, cultures exposed to ciprofloxacin or moxifloxacin displayed a strong dependency on RecA for adaptation, compared to the ones exposed to enrofloxacin or levofloxacin. Genomic analysis further uncovered distinct mutation profiles among the fluoroquinolone-exposed cultures, with notable differences in mutation patterns between the different antibiotics. These findings emphasize the importance of considering the specific fluoroquinolone used in studies of bacterial adaptation and the necessity of studying the dissimilarities between antibiotic compounds belonging to the same class.

## 2. Material and methods

### Strains, growth conditions, and antimicrobial agents

*Escherichia coli* strain MG1655 was used as the wild-type strain. The single gene knockout mutant JW2669 (ΔrecA636::kan) was chosen from the KEIO collection and ordered from Horizon Discovery Ltd. The knockout mutant contained a resistance cassette with flanking FLP recognition elements which was removed using the pCP20 method before starting the experiments (12,13). Liquid or solid lysogeny broth (LB) (10 g/L NaCl) was used to culture the bacteria. Both strains were grown with a starting OD_600_ of 0.1 at 37°C and 200 rpm overnight. For weekend incubations the starting OD_600_ was reduced to 0.01 and the incubation temperature set to 30°C. Fluoroquinolone stocks were purchased from Sigma Aldrich and stored after preparation (10 mM) and filter sterilization either at 4°C for up to one week or at - 20°C for a maximum of a month.

### Minimum inhibitory concentrations (MIC)

MICs were measured twice a week in duplicate for each strain by the broth microdilution method described in (14). In brief, each strain was inoculated into the wells with a starting OD_600_ of 0.05. Readings were performed in a microtiter plate reader every 10 min with 5 min shaking intervals. The MIC was set to the minimal concentration of the antibiotic which inhibited bacterial growth to an OD<0.2 after 24 h incubation.

### Evolution experiment

Evolution experiments were performed as described previously (15). At the beginning of the experiment, the MICs of both strains were determined. The starting concentration for each combination of strain and antibiotic equalled ¼ of the MIC for the specific strain and can be found in the supplementary data (Table 4). After overnight incubation, the OD_600_ was measured. If the value was above 65% of the OD_600_ of the previous culture, the culture was considered to have adapted to the antibiotic, and the antimicrobial concentration was increased 2-fold in fresh medium for the next incubation. If the yield was below this threshold, the culture was transferred to fresh medium with the same antimicrobial concentration as used the previous day. In parallel, each bacterial strain was cultured without antibiotic exposure serving as biological controls. Three biological replicates were performed for each condition. The experiments were stopped after 30 transfers.

### DNA isolation and whole genome sequencing

The DNA of each culture was isolated at the start, after 14 transfers, and at the end of the evolution experiment (30 transfers). After pelleting, phenol-chloroform-isoamyl alcohol (PCI) (Carl Roth) isolation was used to extract the DNA from the whole population. Genomic DNA libraries were created according to the manufacturer’s protocol using NEBNext Ultra II FS DNA Library Prep kit for Illumina (New England BioLabs) in combination with NEBNext multiplex oligos for Illumina (96 Unique Dual Index Primer Pairs; New England BioLabs). The samples were sequenced using a NextSeq 500/550 Mid Output V2.5 kit (Illumina).

### Bioinformatic analysis

The bioinformatics analysis pipeline involved several steps to assess the quality of the sequencing data, trim low-quality reads, align the reads to the reference genome, and perform variant calling. First, the quality of the raw reads was evaluated using FastQC v0.11.9 and MultiQC v1.11. Next, TrimGalore! was used to remove low-quality bases and adapter sequences from the reads. The TrimGalore! analysis was performed using v0.6.7 and cutadapt v1.18 with paired-end trimming. The adapter sequence 5’ – AGATCGGAAGAGC - 3’ corresponding to Illumina TruSeq libraries was auto-detected. The maximum allowed error rate was set to the default value of 0.1 during adapter trimming. Additionally, a minimum required adapter overlap of 1 base pair and a minimum sequence length of 20 base pairs for both reads were enforced to ensure stringent trimming criteria. The trimmed reads were then aligned to the reference genome using Snippy v4.6.0, a tool specifically designed for bacterial variant calling from short-read sequencing data. As reference genomes, U00096 *Escherichia coli* K-12 sub-strain MG1655, and NZ_CP009273 *Escherichia coli* BW25113, also derived from K-12, were used. The sequences were obtained from the NCBI database. Finally, variant calling was performed using Snippy to identify single nucleotide polymorphisms (SNPs) and small insertions/deletions (indels) in comparison to the reference genome. The final data analysis and visualization were performed in Microsoft Excel, R, and Cytoscape. Upon comparison with the reference genomes U00096 and NZ_CP009273, we found that both our susceptible *E. coli* strains exhibited notable genetic variations that were considered for further downstream analysis.

## 3. Results

### Dependency on *recA* for resistance evolution differs between fluoroquinolones

Comparing the onset of resistance towards the tested antibiotics, we could observe a clear pattern in both strains. Both MG1655 and Δ*recA* acquired resistance the fastest when exposed to moxifloxacin. The slowest resistance development was observed during ciprofloxacin exposure. An overview of the number of transfers it took to reach the EUCAST breakpoints can be found in Table 1.

**Table 1.** Comparison average transfers until clinical resistance.

| Fluoroquinolone | MG1655 | $\Delta recA$ |
| --- | --- | --- |
| Ciprofloxacin | 7.7±0.6 | 17±5.3 |
| Enrofloxacin | 5.3*±0.6; 9**±1 | 14.3*±2.5; 26**(*)±2.8 |
| Levofloxacin | 6.7±1.2 | 11.3±3.2 |
| Moxifloxacin | 3.3±0.6 | 7.7±2.5 |

\*Horses. \*\*Poultry. (\*) One replicate could not reach the resistance breakpoint until the experiment ended.

**Table 2.**
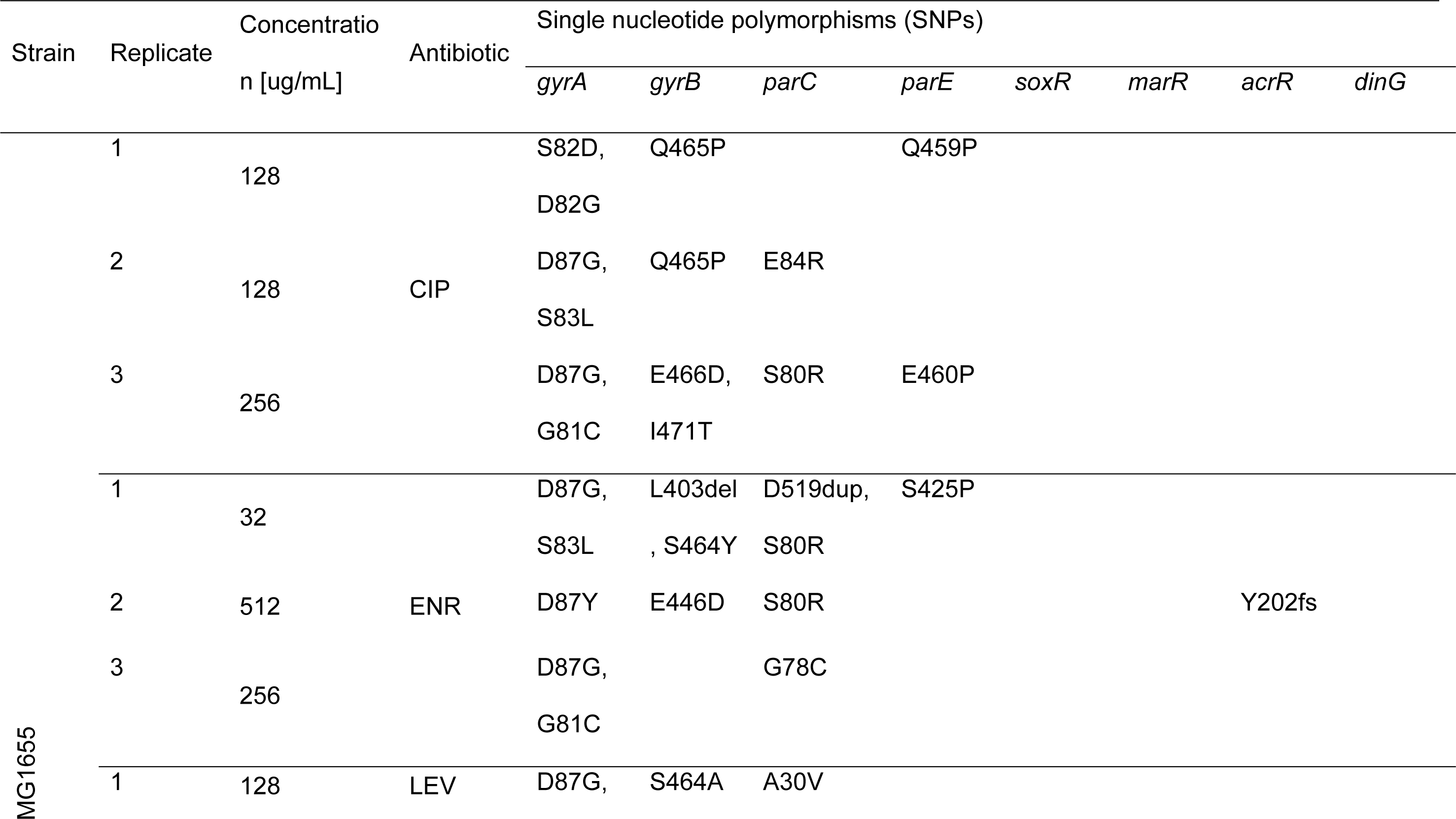
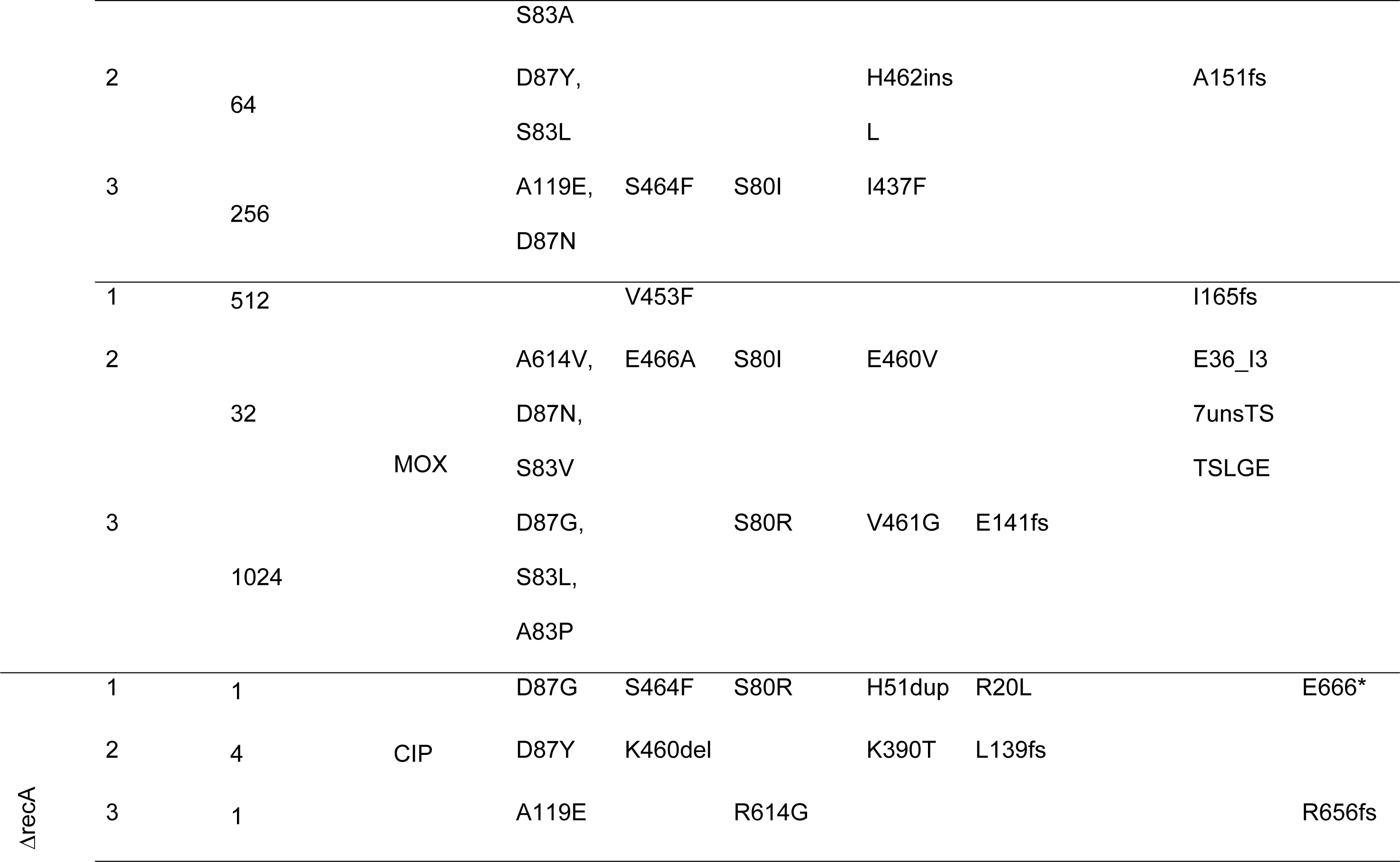
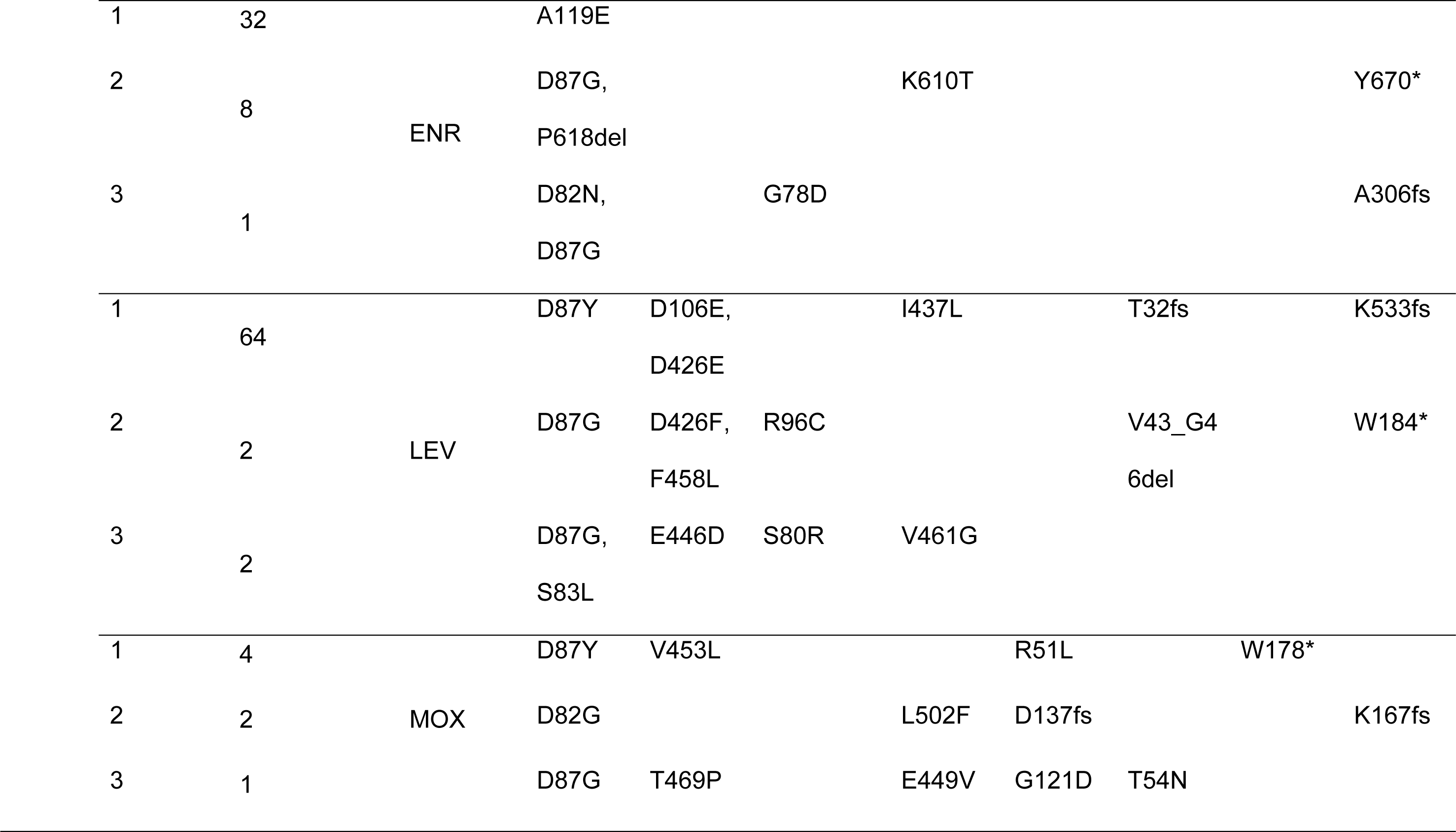
Overview SNPs.

**Table 3.**
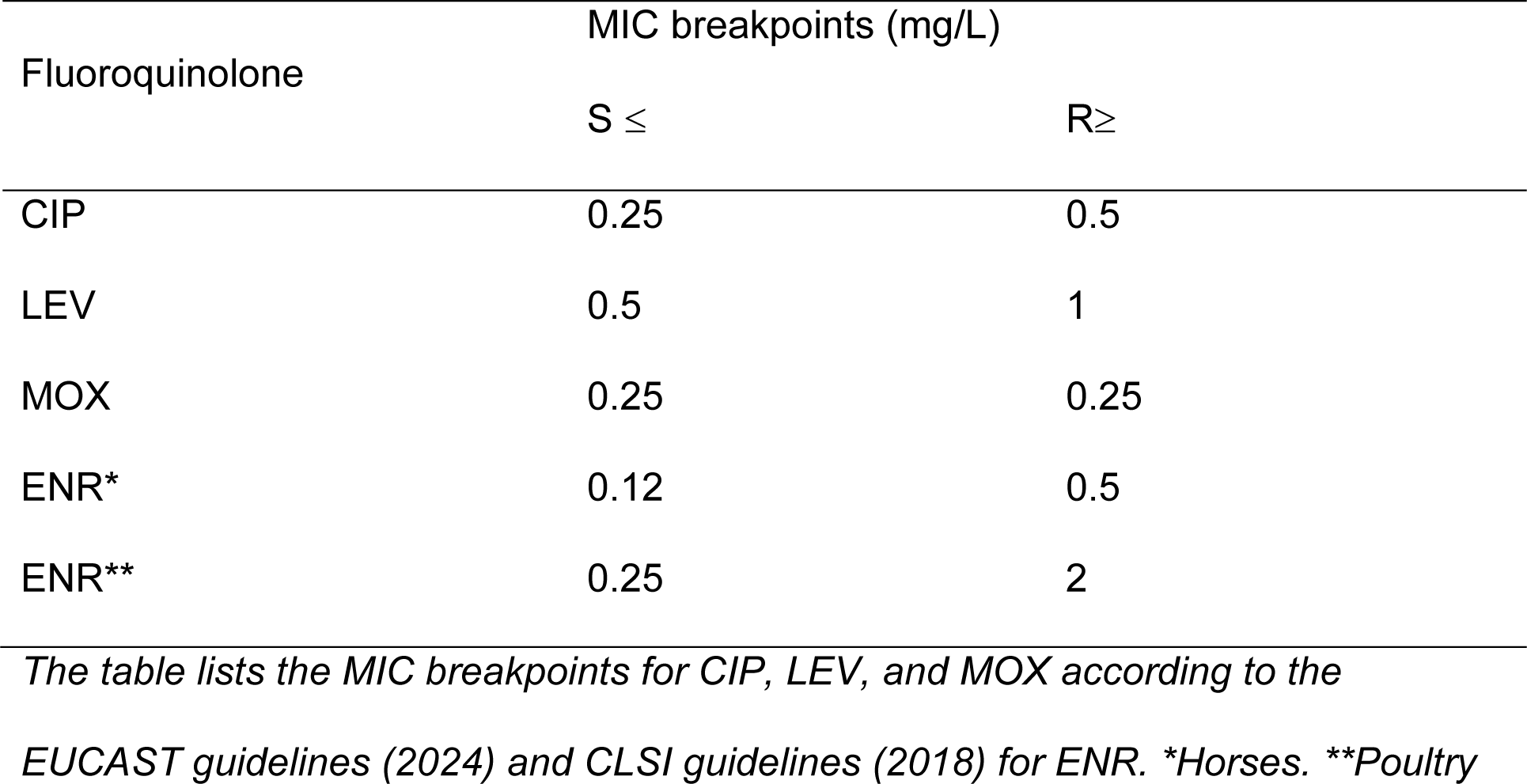
MIC breakpoints.

Relative to the wild-type strain, the knockout mutant (Δ*recA*) displayed a higher inconsistency in adaptation rates to the tested fluoroquinolones between the replicates. In general, the Δ*recA* mutant showed an unpredictable response to antibiotic exposure, often leading to the loss of cultures after incubation in antibiotic concentrations they already adapted to before. The starting concentrations for the evolution experiment with the knockout mutant were lower, reflective of the decreased minimum inhibitory concentration (MIC) of the naive mutant observed during initial MIC testing. The starting concentrations, which equalled ¼ of the starting MICs can be found in Table 4 of the supplementary data.

**Table 4.** Starting concentrations evolution experiment.

| Antibiotic | Antibiotic concentration (mg/L) |  |
| --- | --- | --- |
|  | MG1655 | ΔrecA |
| CIP | 0.015625 | 0.00390625 |
| ENR | 0.03125 | 0.00390625 |
| LEV | 0.03125 | 0.0078125 |
| MOX | 0.0625 | 0.01562 |

*The table compares the starting concentrations for the evolution experiment for each strain and antibiotic. Starting concentrations were ¼ of the initial MIC of the susceptible, non-exposed strain.*

Knocking out *recA* led to a distinct impairment of adaptation to all tested fluoroquinolones (Figure 1). However, there were notable differences between technical replicates, and there also seemed to be a distinction between the different fluoroquinolones. The strongest dissimilarity between Δ*recA* and the wild-type could be observed upon moxifloxacin exposure. While the wild-type could adapt to moxifloxacin concentrations of 32, 512, and 1024 ug/mL, Δ*recA* could only be exposed to a maximum moxifloxacin concentration of 1, 2, or 4 ug/mL, respectively. Similar patterns were found in the ciprofloxacin cultures. Two of the three Δ*recA* replicates could not adapt to ciprofloxacin concentrations above 1 ug/mL during the tested time frame. This pattern supports the hypothesis that targeting the SOS response in bacteria could be a valuable approach to prevent the development of antimicrobial resistance. However, one of the ciprofloxacin-exposed Δ*recA* replicates reached the EUCAST threshold of 0.5 ug/mL after 11 transfers and could survive a final concentration of 4 ug/mL. Yet, this end concentration was considerably lower than the final concentrations of the wild-type replicates, i.e. 128 ug/mL and 256 ug/mL of ciprofloxacin. A detailed account of the concentrations at each transfer cycle is available in the supplementary data (Table 5 and Table 6).

**Figure 1.**
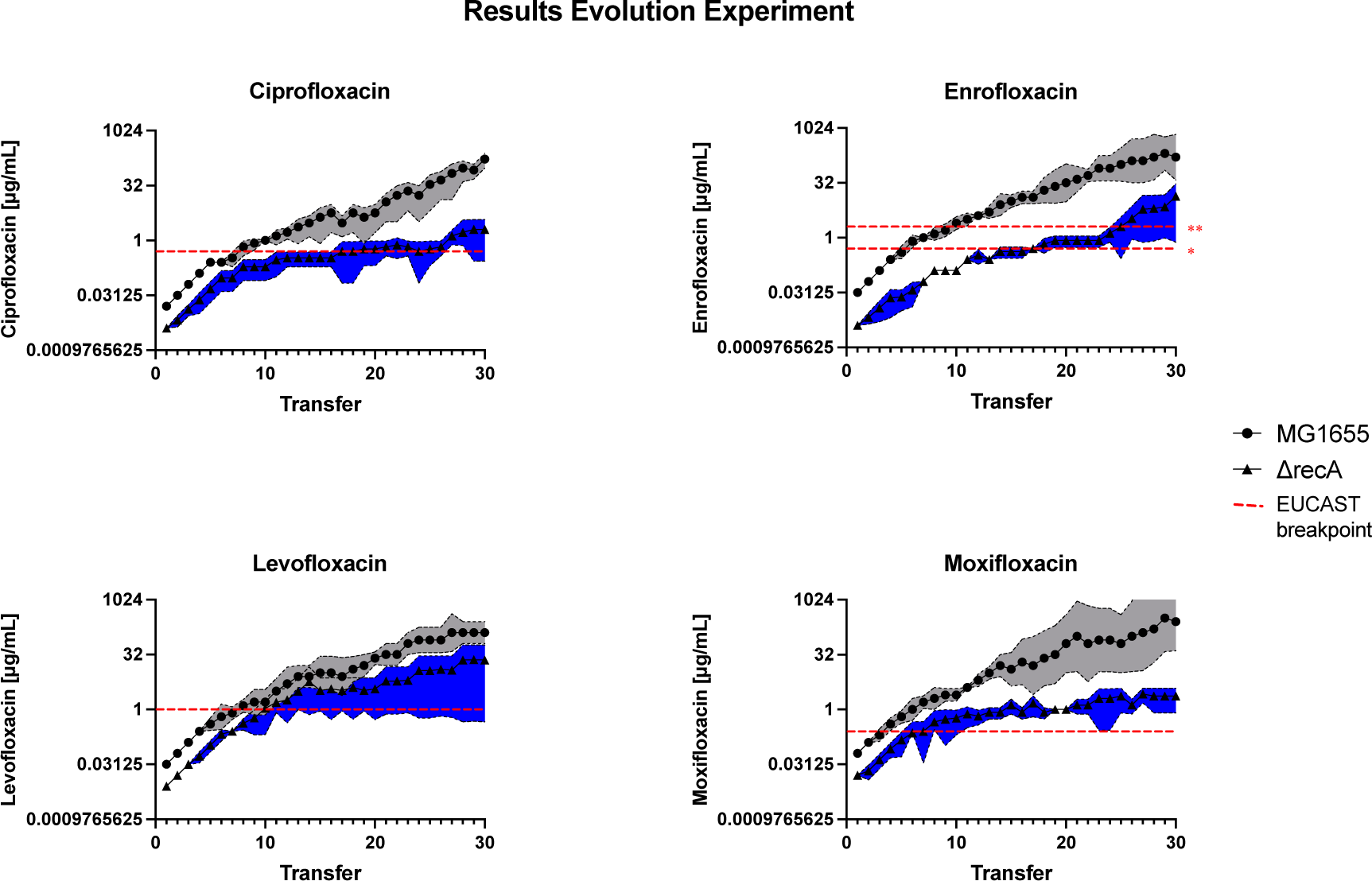
Results Evolution Experiment. Different adaptation patterns of the wild-type and ΔrecA cultures. While the starting concentrations differed between all MG1655 and the corresponding ΔrecA cultures, adaptation rates to levofloxacin and enrofloxacin were comparable. The EUCAST breakpoints can be found in the supplementary data (Table 3). *Horses. **Poultry.

**Table 5.**
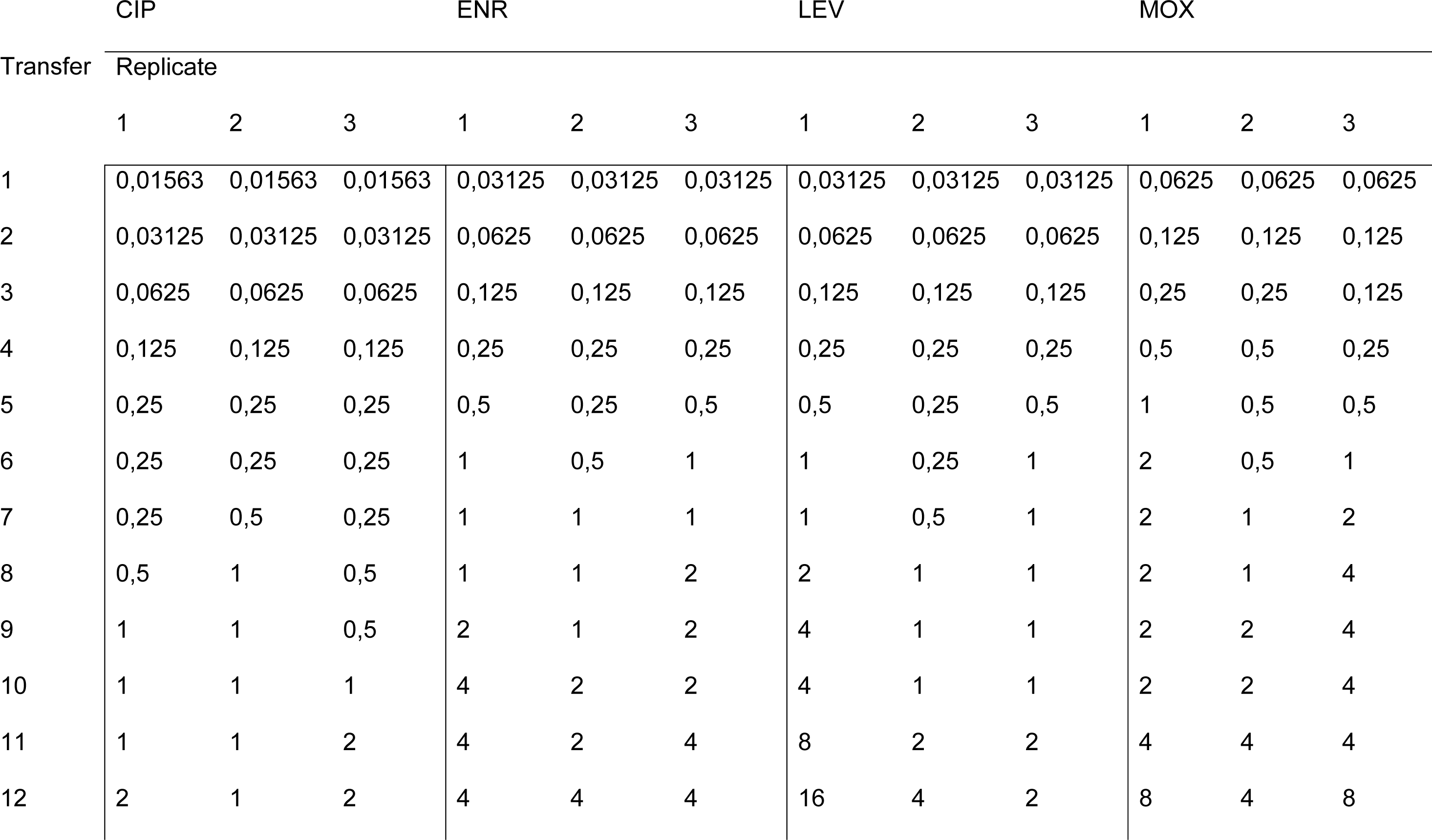
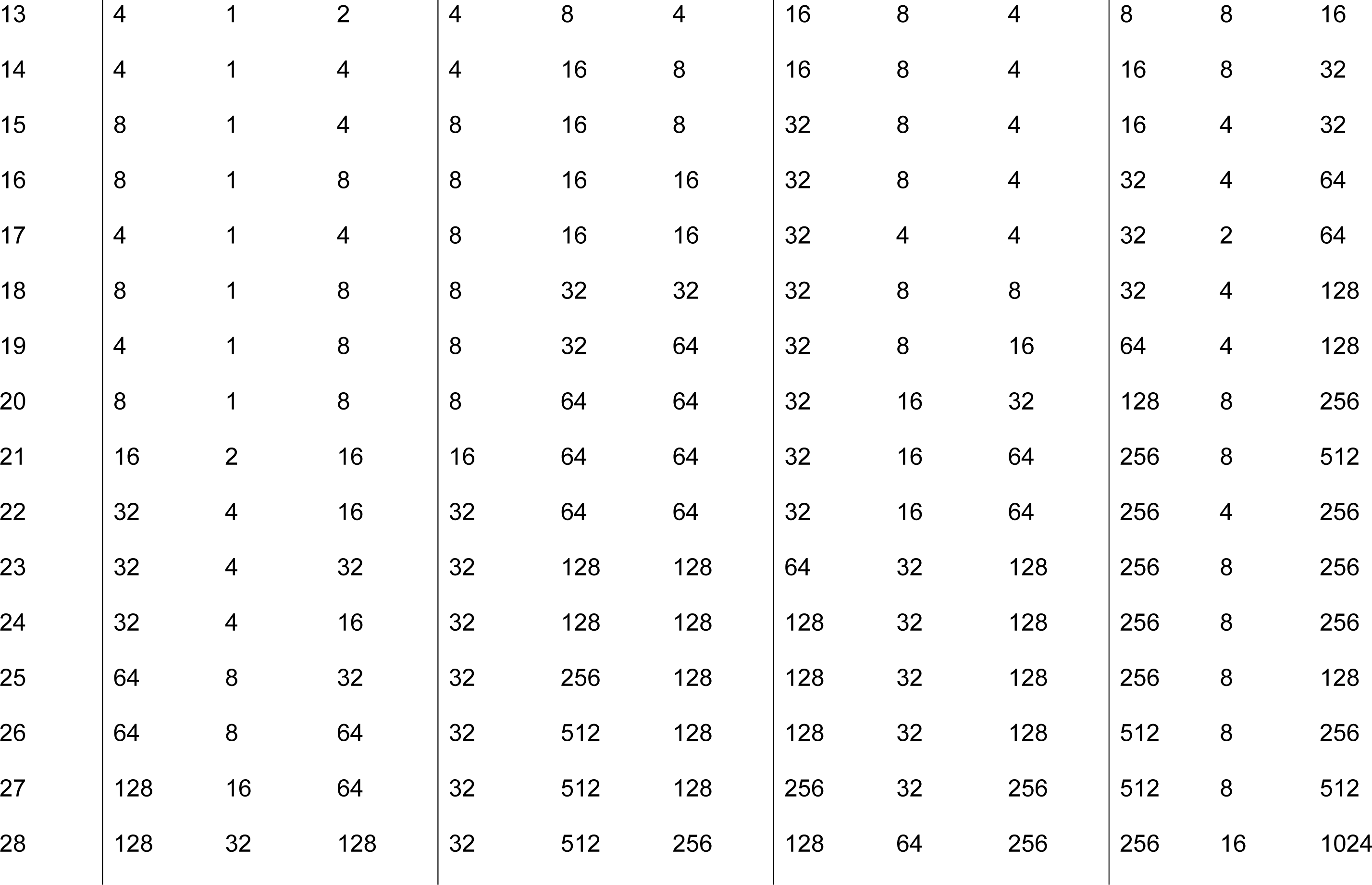
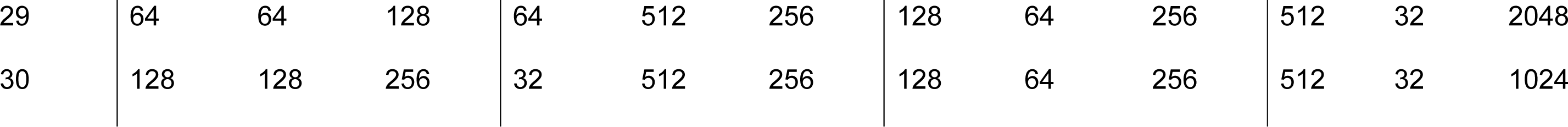
Concentrations evolution experiment. MG1655.

**Table 6.** Concentrations evolution experiment. Knockout mutant.

|  | CIP |  |  | ENR |  |  | LEV |  |  | MOX |  |  |
| --- | --- | --- | --- | --- | --- | --- | --- | --- | --- | --- | --- | --- |
| Transfer | Replicate |  |  |  |  |  |  |  |  |  |  |  |
|  | 1 | 2 | 3 | 1 | 2 | 3 | 1 | 2 | 3 | 1 | 2 | 3 |
| 1 | 0,00391 | 0,00391 | 0,00391 | 0,00391 | 0,00391 | 0,00391 | 0,00781 | 0,00781 | 0,00781 | 0,01563 | 0,01563 | 0,01563 |
| 2 | 0,00781 | 0,00781 | 0,00390625 | 0,00781 | 0,00391 | 0,00781 | 0,015625 | 0,015625 | 0,015625 | 0,01563 | 0,03125 | 0,01563 |
| 3 | 0,015625 | 0,015625 | 0,00781 | 0,015625 | 0,00391 | 0,015625 | 0,03125 | 0,03125 | 0,03125 | 0,03125 | 0,0625 | 0,03125 |
| 4 | 0,03125 | 0,03125 | 0,00781 | 0,03125 | 0,00391 | 0,03125 | 0,0625 | 0,03125 | 0,0625 | 0,0625 | 0,125 | 0,0625 |
| 5 | 0,0625 | 0,0625 | 0,015625 | 0,03125 | 0,00781 | 0,03125 | 0,125 | 0,0625 | 0,125 | 0,125 | 0,25 | 0,0625 |
| 6 | 0,125 | 0,125 | 0,03125 | 0,03125 | 0,015625 | 0,0625 | 0,25 | 0,125 | 0,25 | 0,125 | 0,5 | 0,0625 |
| 7 | 0,125 | 0,125 | 0,03125 | 0,0625 | 0,0625 | 0,0625 | 0,25 | 0,25 | 0,25 | 0,125 | 0,5 | 0,125 |
| 8 | 0,25 | 0,25 | 0,0625 | 0,125 | 0,125 | 0,125 | 0,5 | 0,5 | 0,25 | 0,25 | 1 | 0,125 |
| 9 | 0,25 | 0,25 | 0,0625 | 0,125 | 0,125 | 0,125 | 1 | 0,5 | 0,25 | 0,5 | 1 | 0,125 |
| 10 | 0,25 | 0,25 | 0,0625 | 0,125 | 0,125 | 0,125 | 2 | 1 | 0,25 | 0,5 | 1 | 0,25 |
| 11 | 0,25 | 0,5 | 0,125 | 0,25 | 0,25 | 0,25 | 4 | 0,5 | 0,25 | 1 | 1 | 0,25 |
| 12 | 0,25 | 0,5 | 0,25 | 0,25 | 0,25 | 0,5 | 4 | 1 | 0,5 | 0,5 | 1 | 0,5 |
| 13 | 0,25 | 0,5 | 0,25 | 0,25 | 0,25 | 0,25 | 8 | 1 | 0,5 | 1 | 1 | 0,5 |
| 14 | 0,25 | 0,5 | 0,25 | 0,5 | 0,25 | 0,5 | 16 | 1 | 0,5 | 1 | 1 | 0,5 |
| 15 | 0,25 | 0,5 | 0,25 | 0,5 | 0,25 | 0,5 | 8 | 1 | 1 | 2 | 1 | 1 |
| 16 | 0,25 | 0,5 | 0,25 | 0,5 | 0,25 | 0,5 | 8 | 2 | 1 | 1 | 0,5 | 1 |
| 17 | 0,25 | 1 | 0,25 | 0,5 | 0,5 | 0,5 | 8 | 1 | 1 | 2 | 0,5 | 2 |
| 18 | 0,25 | 1 | 0,25 | 0,5 | 1 | 0,5 | 8 | 2 | 2 | 1 | 1 | 0,5 |
| 19 | 0,25 | 1 | 0,5 | 1 | 1 | 0,5 | 8 | 1 | 1 | 1 | 1 | 1 |
| 20 | 0,25 | 1 | 0,5 | 1 | 1 | 0,5 | 8 | 1 | 2 | 1 | 1 | 1 |
| 21 | 0,5 | 1 | 0,5 | 1 | 1 | 0,5 | 16 | 1 | 1 | 2 | 1 | 1 |
| 22 | 0,25 | 1 | 1 | 1 | 1 | 0,5 | 16 | 1 | 1 | 2 | 1 | 1 |
| 23 | 0,5 | 1 | 0,5 | 1 | 1 | 0,5 | 16 | 1 | 2 | 4 | 1 | 1 |
| 24 | 0,25 | 1 | 0,25 | 2 | 1 | 1 | 32 | 2 | 1 | 4 | 1 | 1 |
| 25 | 0,5 | 1 | 0,25 | 4 | 1 | 1 | 32 | 1 | 2 | 4 | 1 | 2 |
| 26 | 0,5 | 1 | 0,5 | 8 | 1 | 1 | 32 | 2 | 4 | 2 | 1 | 1 |
| 27 | 1 | 2 | 1 | 16 | 1 | 1 | 32 | 2 | 2 | 4 | 2 | 2 |
| 28 | 0,5 | 4 | 0,5 | 16 | 2 | 1 | 64 | 2 | 2 | 4 | 2 | 1 |
| 29 | 1 | 4 | 1 | 16 | 4 | 1 | 64 | 4 | 2 | 4 | 2 | 1 |
| 30 | 1 | 4 | 1 | 32 | 8 | 1 | 64 | 2 | 2 | 4 | 2 | 1 |

### Different fluoroquinolones result in distinct mutational patterns

To study the influence of the four tested fluoroquinolones on the mutational trajectories of the wild-type and the mutant, the genome of every culture was sequenced before the evolution experiment as well as after 14 and 30 transfer cycles.

MG1655 and Δ*recA* both acquired mutations that have commonly been described in fluoroquinolone-resistant strains, i.e. in the genes *gyrA*, *gyrB*, *parC* and *parE* (5,16,17). Both strains also developed mutations that have been described to accelerate antibiotic resistance (*acrR*, and *marR*) (5,17,18). Mutations in these genes were overall the most prevalent and abundant, along with mutations in *soxR* and *dinG*. Both strains gained mutations in genes that were specific for either the wild-type or the knockout mutant. Genes that were only mutated in one replicate were ignored in this analysis. After this filtering, the specific mutations for the strains occurred in *rpoC*, *mprA*, *eptB*, and *rrlF* for MG1655, and *dinG* and *rbsR* for Δ*recA*, respectively. Mutations in *acrR*, *gyrA*, *gyrB*, and *marR* occurred in both strains before transfer 14. Mutations in *parC* and *parE* were acquired by single replicates before transfer 14 but were more abundant at the end of the experiment (Figure 2). This early appearance of *gyrA* mutations and the later onset of *parC* mutations is coherent with earlier observations (19,20).

**Figure 2.**
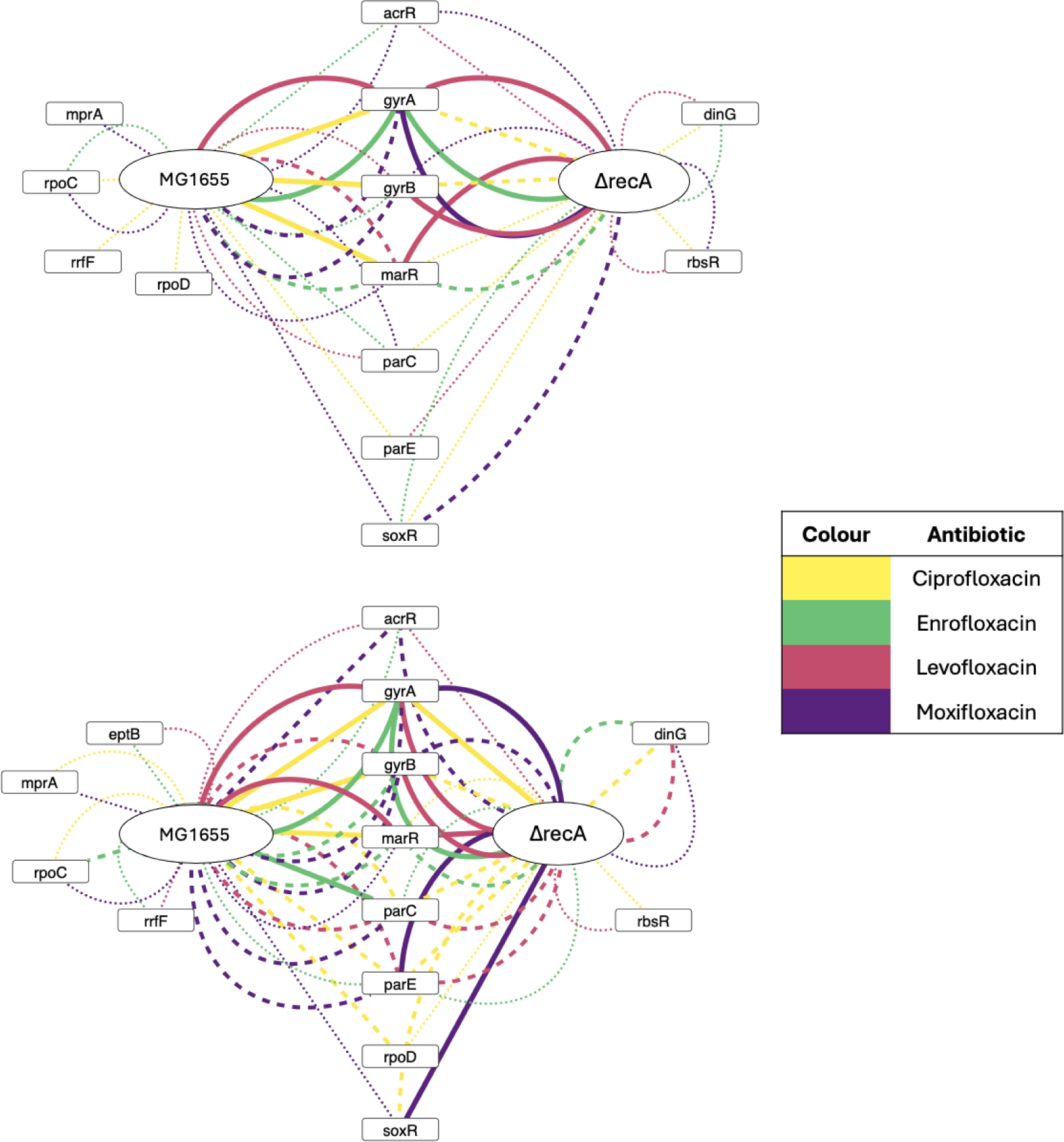
Transfer 14 (upper) and Transfer 30 (lower). The weight of the edges corresponds to the number of replicates with a mutation in each gene. A thin dotted line reflects one replicate, a thick dotted line two, and a thick continuous line equals three replicates.

Distinct patterns could be found at the end of the experiment when examining the individual antibiotic-associated mutations (see network in Figure 2 and Figure 3). Whereas no *acrR* mutations could be observed in any of the ciprofloxacin-exposed strains, it was the only fluoroquinolone that resulted in *rpoD* mutations. Next to moxifloxacin, it was also the only fluoroquinolone associated with *soxR* mutations. Overall, *soxR* mutations were more abundant in the knockout strains than in the wild-type. Mutations in *gyrB* were the least common in enrofloxacin-exposed strains, with no occurrence in any of the knockout replicates. In contrast, all levofloxacin-exposed Δ*recA* replicates had at least a single mutation in *gyrB*. Exposure to levofloxacin led to mutations in *marR* in all cultures, whereas exposure to moxifloxacin resulted in a mutation in this gene for only one of the wild-type replicates. Adaptation to moxifloxacin was possible for all Δ*recA* replicates without mutations in *parC*. Noteworthily, one of the moxifloxacin-exposed wild-type replicates could adapt to a concentration of 512 ug/mL without any *gyrA*, *parC,* or *parE* mutation.

**Figure 3.**
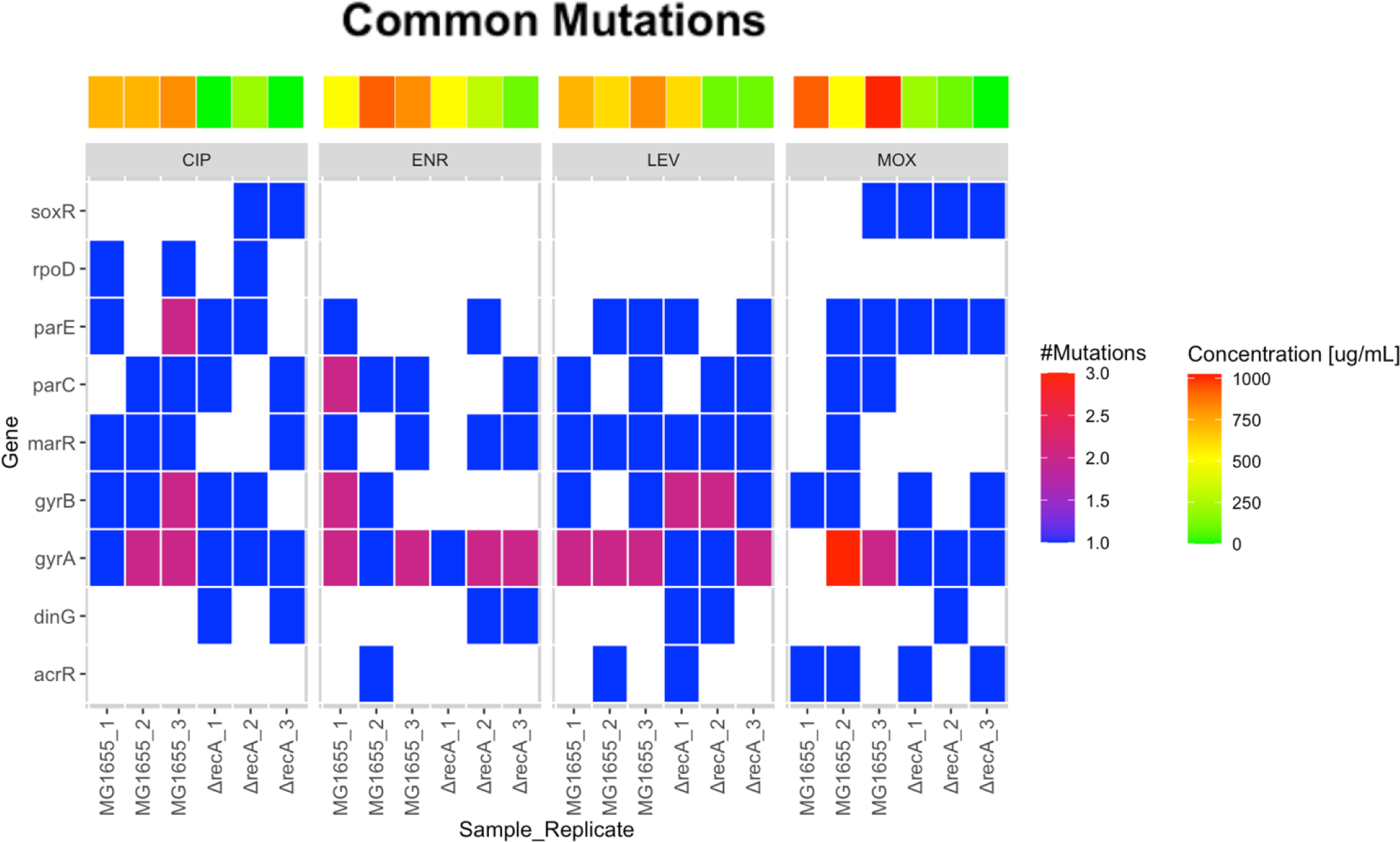
Heatmap Common Mutations. Most common mutations found in the wild-type and knockout mutant after 30 transfers.

Our evolved strains showed a higher prevalence of D87 than S83 mutations in the DNA gyrase subunit A (see Table 2). 80% of the strains had a D87 mutation while only 30% of the evolved strains had a mutation in S83. It is noteworthy that this pattern was pronounced in the knockout mutant – while 75% of the evolved strains carried a D87 mutation, only a single strain carried an S83 mutation at the end of the evolution experiment. In addition, 75% of the knockout strains had only a single mutation in *gyrA*, whereas 83% of the wild-type had two mutations in the DNA gyrase.

## 4. Discussion

The observed patterns of mutations accompanying the development of resistance differed strongly between the four fluoroquinolones used in this study. Examining the adaptation in response to fluoroquinolone exposure, as well as its dependency on the SOS response, revealed intriguing dynamics. Notably, the Δ*recA* knockout and wild-type strains displayed the fastest development of clinical levels of resistance following moxifloxacin exposure, while the slowest adaptation rate could be observed during ciprofloxacin exposure. Furthermore, our analysis indicates that moxifloxacin- and ciprofloxacin-exposed strains exhibited the strongest reliance on RecA for their adaptation, in contrast to enrofloxacin- and levofloxacin-exposed cultures. The success of impairing the SOS response as a potential resistance evolution inhibition strategy seems to depend not only on biological variation but also on the specific antibiotic used. The difference in SOS response induction upon exposure between different fluoroquinolones is not commonly recognized. On the contrary, it is generally assumed that the cell’s response to the different antibiotics within a class is interchangeable (21–27).

While the differences in chemical properties of each fluoroquinolone seem apparent, the impact of these differences on the cellular uptake and target binding remains unclear. A computational quinolone-gyrase binding study indicated different binding affinities of moxifloxacin, ciprofloxacin, and levofloxacin to the gyrase (28). While the commonly described S83L mutation in *gyrA* seems to cause high-level resistance to ciprofloxacin and levofloxacin, its impact on moxifloxacin resistance seems less important due to improved binding of this drug to the GyrA protein (28). Furthermore, the study indicated that D87 and R121 mutations are more critical binding sides for ciprofloxacin, levofloxacin, and moxifloxacin than S83 (28). The evolved strains indeed showed a higher prevalence of D87 than S83 mutations in the DNA gyrase (see Table 2), indicating varying binding affinities of the different fluoroquinolones and thus diverging impact on resistance evolution.

The genomic analysis of the resistant cultures further unveiled distinct mutation profiles by comparing the effects of different fluoroquinolones to each other. We less frequently observed mutations in *marR* in moxifloxacin-exposed strains in comparison to the other antibiotic-exposed strains, especially levofloxacin, which induced *marR* mutations in all cultures. The *mar* regulon has not only been proposed to be involved in reduced drug uptake and AcrAB efflux pump regulation, but also the repair of quinolone-induced DNA damage (29–31). It is noteworthy, however, that SoxS expression changes, partially regulated by *soxR*, have been suggested as an alternative to *mar* overexpression as an instrument for developing resistance (31). Our analysis indicates that moxifloxacin exposure results in a higher prevalence of *acrR* and *soxR* mutations than the other fluoroquinolones, implying a change in AcrAB efflux pump expression in the moxifloxacin-resistant strains via these regulons (32). Nevertheless, for moxifloxacin-resistant cultures, efflux might not be the main resistance mechanism due to the chemical complexity and hydrophobic properties of the antibiotic (33).

The biological variability observed among our replicates could be partially attributed to specific single nucleotide polymorphisms (SNPs). One of the three moxifloxacin-exposed wild-type replicates acquired a S80I mutation in *parC*. This replicate was able to reach a concentration of only 32 ug/mL, in comparison to the corresponding replicates which reached 1024 and 512 ug/mL. This amino acid substitution has been shown to increase mutation rates in *E. coli* under moxifloxacin exposure 1300-fold (34). It could lead to a reduction in population fitness by the accumulation of deleterious mutations. This lower fitness can be further reduced by passage through strong bottlenecks (Muller’s ratchet) (35–37) or by driving out of the most adapted genotype, because the population gets overwhelmed with the mutational backflow (38). A higher mutation rate is beneficial for resistance acquired by several mutations, such as is the case for high-level fluoroquinolone resistance (39,40). However, it is unclear at which mutation rate the benefit becomes a disadvantage. Indeed, in natural *E. coli* isolates, weak mutators were found to have the highest antibiotic resistance (41).

It is noteworthy that Schedletzky and co-workers described different mutation rates due to the S80I mutation in *parC*, depending on the antibiotic used (34). Ciprofloxacin exposure caused a lower mutation rate increase (40-fold in comparison to the 1300-fold increase during moxifloxacin exposure), possibly affecting the population’s fitness less. This finding seems consistent with our observed phenotypes. While the wild-type that gained this mutation could only reach a concentration of 32 ug/mL under moxifloxacin exposure, another one of the wild-type replicates with this substitution could adapt to a levofloxacin concentration of 256 ug/mL, the highest one observed among the replicates. Furthermore, Moxifloxacin has been described to have a higher impact on the growth rate at sub-MIC levels than ciprofloxacin (42), yet it has been reported that it induces the SOS response to a lesser degree than ciprofloxacin (43). This observation can be explained by the increase in SOS response at higher metabolic rates caused by the higher ROS levels in cells that metabolize fast (44).

In addition to studying individual mutations to explain certain phenomena, it is essential to analyse the full genomic profile of an evolved strain to understand the potential synergetic effects of mutations on the observed phenotype. All strains did adapt to a certain extent to the fluoroquinolones, whether they possessed a functional *recA* gene or not. Whether this adaptation occurred due to RecA-independent activation of the SOS response or due to activation of other pathways remains unclear. While ciprofloxacin exposure seems to cause downregulation of the mismatch repair system (45) inactivation of the mismatch repair system alone is not sufficient to acquire ciprofloxacin-resistance-inducing mutations if the SOS response is impaired (46).

It has been proposed that parts of the SOS response can be activated independently of the LexA/RecA-regulon upon beta-lactam exposure and that the resistance-inducing mutations are caused by upregulation of *dinB*, an error-prone DNA polymerase that is part of the SOS response (47). However, a *dinB* knockout mutant can adapt to amoxicillin, and enrofloxacin, at almost the same rate as a wild-type *E. coli* (Teichmann *et al*., in preparation), indicating that the DNA polymerase is not essential for adaptation. Alternative mechanisms like adaptive amplification and amplification-mutagenesis have also been suggested (48).

Our analysis highlights that the patterns of adaptation and genetic alterations depend on the fluoroquinolone used, underscoring potential consequences for healthcare settings. Therefore, studies primarily using, e.g., ciprofloxacin may not fully capture the extent of responses elicited by the class of fluoroquinolones. Moreover, the identification of specific mutations associated with fluoroquinolone resistance emphasizes the importance of considering the diversity of fluoroquinolone responses in clinical practice. Failure to account for this variability may have significant implications for antibiotic stewardship, treatment efficacy, and the emergence of antibiotic-resistant bacteria.

## 6. Supplementary data

## Literature

1. Brown SA. Fluoroquinolones in animal health. J Vet Pharmacol Ther. 1996;19(1):1–14.

2. Wolfson JS, Hooper DC. Fluoroquinolone antimicrobial agents. Clin Microbiol Rev. 1989;2(4):378–424.

3. World Health Organization. 2021 AWaRe classification [Internet]. 2021. Available from: https://www.who.int/publications/i/item/2021-aware-classification

4. Hooper DC. Mechanisms of action of antimicrobials: Focus on fluoroquinolones. Clinical Infectious Diseases. 2001;32(SUPPL. 1):9–15.

5. Hooper DC. Emerging mechanisms of fluoroquinolone resistance. Emerg Infect Dis. 2001;7(2):337–41.

6. Cirz RT, Chin JK, Andes DR, De Crécy-Lagard V, Craig WA, Romesberg FE. Inhibition of mutation and combating the evolution of antibiotic resistance. PLoS Biol. 2005;3(6):1024–33.

7. Yakimov A, Bakhlanova I, Baitin D. Targeting evolution of antibiotic resistance by SOS response inhibition. Comput Struct Biotechnol J. 2021;19:777–83.

8. Maslowska KH, Makiela-Dzbenska K, Fijalkowska IJ. The SOS system: A complex and tightly regulated response to DNA damage. Environ Mol Mutagen. 2019;60(4):368–84.

9. Lusetti SL, Cox MM. The bacterial RecA protein and the recombinational DNA repair of stalled replication forks. Annu Rev Biochem. 2002;71:71–100.

10. Erill I, Campoy S, Barbé J. Aeons of distress: An evolutionary perspective on the bacterial SOS response. FEMS Microbiol Rev. 2007;31(6):637–56.

11. Pérez-Capilla T, Baquero MR, Gómez-Gómez JM, Ionel A, Martín S, Blázquez J. SOS-independent induction of dinB transcription by β-lactam-mediated inhibition of cell wall synthesis in Escherichia coli. J Bacteriol. 2005;187(4):1515–8.

12. Baba T, Ara T, Hasegawa M, Takai Y, Okumura Y, Baba M, et al. Construction of Escherichia coli K-12 in-frame, single-gene knockout mutants: The Keio collection. Mol Syst Biol. 2006 May 16;2.

13. Datsenko KA, Wanner BL. One-step inactivation of chromosomal genes in Escherichia coli K-12 using PCR products. PNAS. 2000;97(12):6640–5.

14. Schuurmans JM, Nuri Hayali AS, Koenders BB, ter Kuile BH. Variations in MIC value caused by differences in experimental protocol. J Microbiol Methods. 2009 Oct;79(1):44–7.

15. Van Der Horst MA, Schuurmans JM, Smid MC, Koenders BB, Ter Kuile BH. De novo acquisition of resistance to three antibiotics by escherichia coli. Microbial Drug Resistance. 2011 Jun 1;17(2):141–7.

16. Redgrave LS, Sutton SB, Webber MA, Piddock LJV. Fluoroquinolone resistance: Mechanisms, impact on bacteria, and role in evolutionary success. Trends Microbiol. 2014;22(8):438–45.

17. Okusu H, Ma D, Nikaido H. AcrAB efflux pump plays a major role in the antibiotic resistance phenotype of Escherichia coli multiple-antibiotic-resistance (Mar) mutants. J Bacteriol. 1996;178(1):306–8.

18. Marcusson LL, Frimodt-Møller N, Hughes D. Interplay in the selection of fluoroquinolone resistance and bacterial fitness. PLoS Pathog. 2009;5(8).

19. Tavío M del M, Vila J, Ruiz J, Ruiz J, Martín-Sánchez AM, Jiménez de Anta MT. Mechanisms involved in the development of resistance to fluoroquinolones in Escherichia coli isolates. Journal of antimicrobial chemotherapy. 1999;44(1):735–42.

20. Händel N, Hoeksema M, Mata MF, Brul S, Ter Kuile BH. Effects of stress, reactive oxygen species, and the SOS response on de novo acquisition of antibiotic resistance in Escherichia coli. Antimicrob Agents Chemother. 2016;60(3):1319–27.

21. Dörr T, Lewis K, Vulić M. SOS response induces persistence to fluoroquinolones in Escherichia coli. PLoS Genet. 2009 Dec;5(12).

22. Critchlow SE, Maxwell A. DNA Cleavage Is Not Required for the Binding of Quinolone Drugs to the DNA Gyrase-DNA Complex. Biochemistry. 1996;35:7387–93.

23. Palù G, Valisena S, Ciarrocchit G, Gattot B, Palumbo M. Quinolone binding to DNA is mediated by magnesium ions. PNAS. 1992;89:9671–5.

24. Da Re S, Garnier F, Guérin E, Campoy S, Denis F, Ploy MC. The SOS response promotes qnrB quinolone-resistance determinant expression. EMBO Rep. 2009;10(8):929–33.

25. Foti JJ, Devadoss B, Winkler JA, Collins JJ, Walker GC. Oxidation of the Guanine Nucleotide Pool Underlies Cell Death by Bactericidal Antibiotics. Science (1979). 2012;336:315–9.

26. Alam MK, Alhhazmi A, Decoteau JF, Luo Y, Geyer CR. RecA Inhibitors Potentiate Antibiotic Activity and Block Evolution of Antibiotic Resistance. Cell Chem Biol. 2016 Mar 17;23(3):381–91.

27. Kohanski MA, Dwyer DJ, Hayete B, Lawrence CA, Collins JJ. A Common Mechanism of Cellular Death Induced by Bactericidal Antibiotics. Cell. 2007 Sep 7;130(5):797–810.

28. Madurga S, Sánchez-Céspedes J, Belda I, Vila J, Giralt E. Mechanism of binding of fluoroquinolones to the quinolone resistance-determining region of DNA gyrase: Towards an understanding of the molecular basis of quinolone resistance. ChemBioChem. 2008 Sep 1;9(13):2081–6.

29. Sharma P, Haycocks JRJ, Middlemiss AD, Kettles RA, Sellars LE, Ricci V, et al. The multiple antibiotic resistance operon of enteric bacteria controls DNA repair and outer membrane integrity. Nat Commun. 2017 Dec 1;8(1).

30. Cohen SP, Mcmurry LM, Hooper DC, Wolfson JS, Levy SB. Cross-Resistance to Fluoroquinolones in Multiple-Antibiotic-Resistant (Mar) Escherichia coli Selected by Tetracycline or Chloramphenicol: Decreased Drug Accumulation Associated with Membrane Changes in Addition to OmpF Reduction. Antimicrob Agents Chemother. 1989;33(8):1318–25.

31. Kern W V, Oethinger M, Jellen-Ritter AS, Levy SB. Non-Target Gene Mutations in the Development of Fluoroquinolone Resistance in Escherichia coli. Antimicrob Agents Chemother. 2000;44(4):814–20.

32. Gerstel A, Beas JZ, Duverger Y, Bouveret E, Barras F, Py B. Oxidative stress antagonizes fluoroquinolone drug sensitivity via the SoxR-SUF Fe-S cluster homeostatic axis. PLoS Genet. 2020 Nov 2;16(11).

33. Pestova E, Millichap JJ, Noskin GA, Peterson LR. Intracellular targets of moxifloxacin: a comparison with other fluoroquinolones. Journal of Antimicrobial Chemotherapy. 2000;45:583–90.

34. Schedletzky H, Wiedemann B, Heisig P. The effect of moxifloxacin on its target topoisomerases from Escherichia coli and Staphylococcus aureus. Journal of Antimicrobial Chemotherapy. 1999;43(Suppl. B):31–7.

35. Springman R, Keller T, Molineux IJ, Bull JJ. Evolution at a high imposed mutation rate: Adaptation obscures the load in phage T7. Genetics. 2010 Jan;184(1):221–32.

36. Kimura M, Maruyama T. The mutational load with epistatic gene interactions in fitness. Genetics. 1966;54:1337–51.

37. Denamur E, Matic I. Evolution of mutation rates in bacteria. Vol. 60, Molecular Microbiology. 2006. p. 820–7.

38. Eigen M, Mccaskill J, Schuster P, Eigen), Mccaskill M, Schuster JS, et al. Molecular Quasi-Species. J Phys Chem. 1988;92:6881–91.

39. Lindgren PK, Karlsson Å, Hughes D. Mutation rate and evolution of fluoroquinolone resistance in Escherichia coli isolates from patients with urinary tract infections. Antimicrob Agents Chemother. 2003 Oct 1;47(10):3222–32.

40. Tenaillon O, Toupance B, Le Nagard H, Taddei F, Godelle B. Mutators, Population Size, Adaptive Landscape and the Adaptation of Asexual Populations of Bacteria. Genetics. 1999;152:485–93.

41. Denamur E, Tenaillon O, Deschamps C, Skurnik D, Ronco E, Gaillard JL, et al. Intermediate mutation frequencies favor evolution of multidrug resistance in Escherichia coli. Genetics. 2005;171(2):825–7.

42. Boswell FJ, Andrews JM, Wise R, Dalhoff A. Bactericidal properties of moxifloxacin and post-antibiotic effect. Journal of Antimicrobial Chemotherapy. 1999;43:43–9.

43. Vestergaard M, Paulander W, Ingmer H. Activation of the SOS response increases the frequency of small colony variants. BMC Res Notes. 2015 Dec 8;8(1).

44. Qi W, Jonker MJ, Katsavelis D, de Leeuw W, Wortel M, ter Kuile BH. The Effect of the Stringent Response and Oxidative Stress Response on Fitness Costs of De Novo Acquisition of Antibiotic Resistance. Int J Mol Sci. 2024 Mar 1;25(5).

45. Machuca J, Recacha E, Briales A, Díaz-de-Alba P, Blazquez J, Pascual álvaro, et al. Cellular response to ciprofloxacin in low-level quinolone-resistant Escherichia coli. Front Microbiol. 2017 Jul 19;8(JUL).

46. Cirz RT, Romesberg FE. Induction and inhibition of ciprofloxacin resistance-conferring mutations in hypermutator bacteria. Antimicrob Agents Chemother. 2006 Jan;50(1):220–5.

47. Pérez-Capilla T, Baquero MR, Gómez-Gómez JM, Ionel A, Martín S, Blázquez J. SOS-independent induction of dinB transcription by β-lactam-mediated inhibition of cell wall synthesis in Escherichia coli. J Bacteriol. 2005 Feb;187(4):1515–8.

48. Hersh MN, Ponder RG, Hastings PJ, Rosenberg SM. Adaptive mutation and amplification in Escherichia coli: Two pathways of genome adaptation under stress. Vol. 155, Research in Microbiology. Elsevier Masson SAS; 2004. p. 352–9.

